# Triparental inheritance in *Dictyostelium*

**DOI:** 10.1101/385708

**Authors:** Gareth Bloomfield, Peggy Paschke, Marina Okamoto, Tim J. Stevens, Hideko Urushihara

**Affiliations:** MRC Laboratory of Molecular Biology, Cambridge, UK; Faculty of Life and Environmental Sciences, University of Tsukuba, Tsukuba, Japan

## Abstract

Sex promotes the recombination and reassortment of genetic material and is prevalent across eukaryotes. In social amoebae sex involves a promiscuous mixing of cytoplasm before zygotes consume the majority of cells. We report here the first genomewide characterisation of meiotic progeny in *Dictyostelium discoideum*. We find that recombination occurs at high frequency in pairwise crosses between all three mating types, despite the absence of the SPO11 enzyme that is normally required to initiate crossover formation. In crosses involving three strains, transient fusions involving more than two gametes frequently lead to triparental inheritance, with recombined nuclear haplotypes inherited from two parents and the mitochondrial genome from a third. Cells that do not contribute genetically to the *Dictyostelium* zygote nucleus thereby have a stake in the next haploid generation. We suggest that this lateral transfer helps to enforce cooperation in this confictual system.

## Introduction

Inheritance of nuclear genes largely follows the Mendelian laws of segregation and independent assortment. Individual eukaryotes are produced either clonally, having a single parent, or sexually with two parents. In contrast, cytoplasmic genes display non-Mendelian patterns of inheritance, and are typically transmitted only from one of the two parents during sex. The uncoupled inheritance of these different sets of genes creates recurrent genetic conflict (1–3). The effects of such conflict are now well understood in diverse animals and plants, but in many less well-studied eukaryotes the effects of sexual conflict are unclear. Social amoebae present intriguing examples of conflict and their modes of inheritance are relatively neglected. These protists proliferate as single cells, feeding on bacteria, but when starved undergo developmental programs involving intimate contacts between conspecific cells. In their asexual cycle, many thousands of social amoebae aggregate to form multicellular structures, ultimately forming fruiting bodies consisting of many individual haploid spore cells atop slender stalks. In most genera, the stalk is composed of dead, vacuolated cells; this leads to the potential for conflict, since any variant that is over-represented among spore cells is at an advantage. The sexual cycle of social amoebae is even more conflictual: as they differentiate, zygotes feed cannibalistically on surrounding cells, including other zygotes as well as haploid amoebae (4). Sex in social amoebae has a number of other unusual features. Several species have more than two mating types (5, 6), which are specified in part by homeodomain-like proteins in *Dictyostelium* (7). Social amoeba gametes are indistinguishable in size (8), and their fusion is reliant on HAP2/GCS1-related proteins (9). Gamete fusion in *Dictyostelium* is unusually promiscuous, with no mechanism preventing multiple gametes from fusing, so that syncytia can form (Fig. 1A) (10–12). In contrast, in most eukaryotes syngamy is controlled to ensure strict genome doubling because accidental polyspermy leads to polyploidy (13, 14). *Dictyostelium* syncytia break apart gradually over the course of several hours giving rise to binucleate cells before nuclear fusion occurs (12); control of nuclear pairing and fusion is not understood in these cells. Zygotes then attract and ingest surrounding cells (Fig. 1B), and grow without undergoing mitosis eventually forming semi-dormant walled diploid cells called macrocysts (Fig. 1C). Meiosis is believed to occur in young macrocysts, where there is ultrastructural evidence for synaptonemal complexes (11, 15). Macrocysts can remain dormant for several weeks before germinating to release haploid amoebae, but the triggers for germination remain unclear so generating progeny in the laboratory is difficult. In one cross in which recombinant *D. discoideum* haploid progeny could be obtained, crossovers on one chromosome were found to be as frequent as in *Saccharomyces cerevisiae*, which has one of the highest crossover frequencies known (16, 17). In retrospect this is surprising: genome sequencing later revealed that although a number of meiosis-specific genes are conserved in social amoebae, the gene encoding SPO11, the transesterase normally required to initiate crossover formation (18) has been lost in this lineage (19). How recombination might occur in the absence of this key enzyme remains mysterious (20).

**Figure 1.**
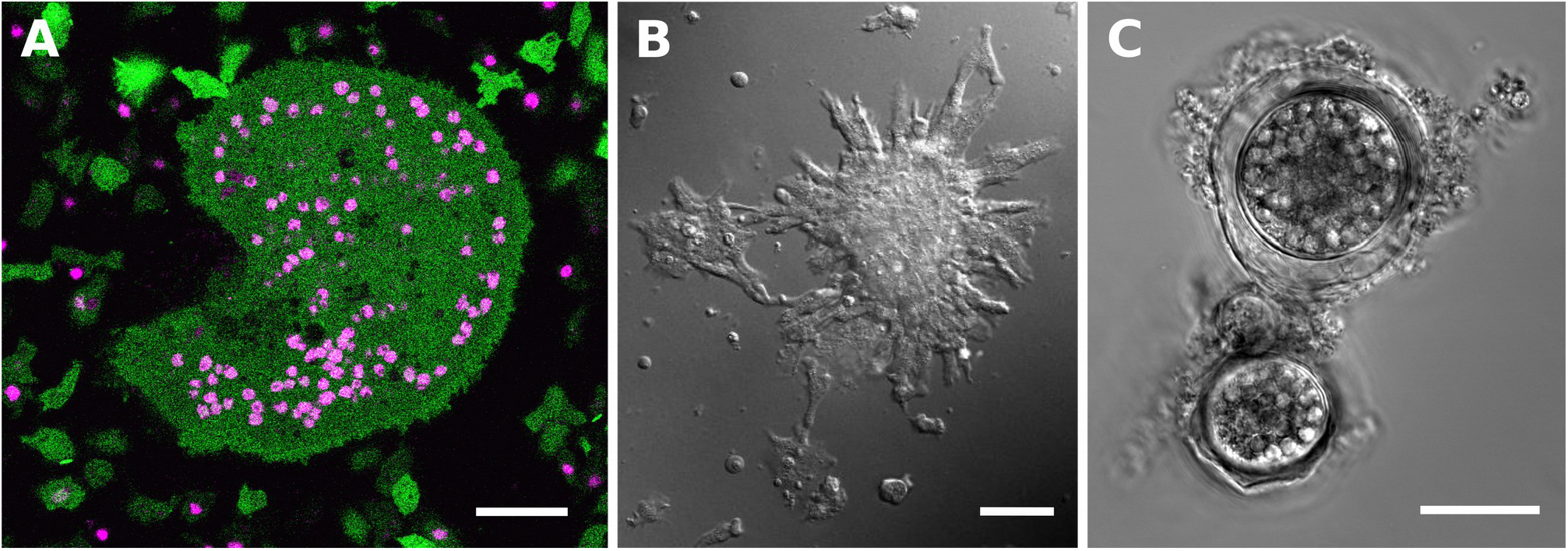
Sexual development in *Dictyostelium*. **A.** A syncytium formed after fusion of a *D. discoideum* strain (HM1558) expressing cytoplasmic GFP with a strain (AX2) expressing RFP-tagged histone 2B, surrounded by smaller fused and unfused amoebae. The cells were made fusion-competent separately, then mixed and shaken together for one hour before incubation in a chamber slide. These cells were imaged seven hours after mixing. **B.** Aggregation around zygotes: cells of strain AC4 were made fusion-competent and incubated in a chamber slide. Several zygotes as well as haploid amoebae aggregate around the large zygote in the centre-right of the field. **C.** Mature macrocysts 15 days after mixing strains HM597 and HM598; cannibalised amoebae are still prominent in food vacuoles inside the walled cysts. Scale bar represents 25 µm in all panels.

## Results and Discussion

To investigate these inheritance patterns in greater detail, we germinated macrocysts from three pairwise crosses between strains of all three *D. discoideum* mating types (HM597, HM598, and WS2162), and sequenced the genomes of haploid progeny clones along with those of their parents. The parental genomes were extensively polymorphic, such that after filtering of putative variants thousands of sites could be used to assess recombination in meiotic progeny across the 34 Mb genome in each cross. The type II parent strain, HM597, is a cycloheximide resistant mutant. The affected *cycA* gene was previously mapped to chromosome 1 (21), but the identity of the affected gene had not been confirmed. We identified a proline-to-serine (P54S) mutation in HM597 in the *rpl36a* gene that encodes the ribosomal protein eL42 and which lies on this chromosome (Fig. S1). Variants at the same amino acid residue in the orthologous protein distinguish cycloheximide-sensitive and -resistant yeast species (22). Initial observations confirmed that *cycA* and the mating-type locus, which lies on chromosome 5, were reassorted in several putative meiotic progeny. Genomewide analysis of progeny from these three crosses revealed that recombination is very frequent, with at least one crossover per chromosome (Figs. 2 and S2). We found that progeny from each macrocyst are identical, implying that only one meiotic product survived in each cyst (Table S1 and Fig. S3). However different macrocysts in a given cross give rise to diverse progeny genotypes (Figs. 3, S2, and S4).

**Figure 2.**
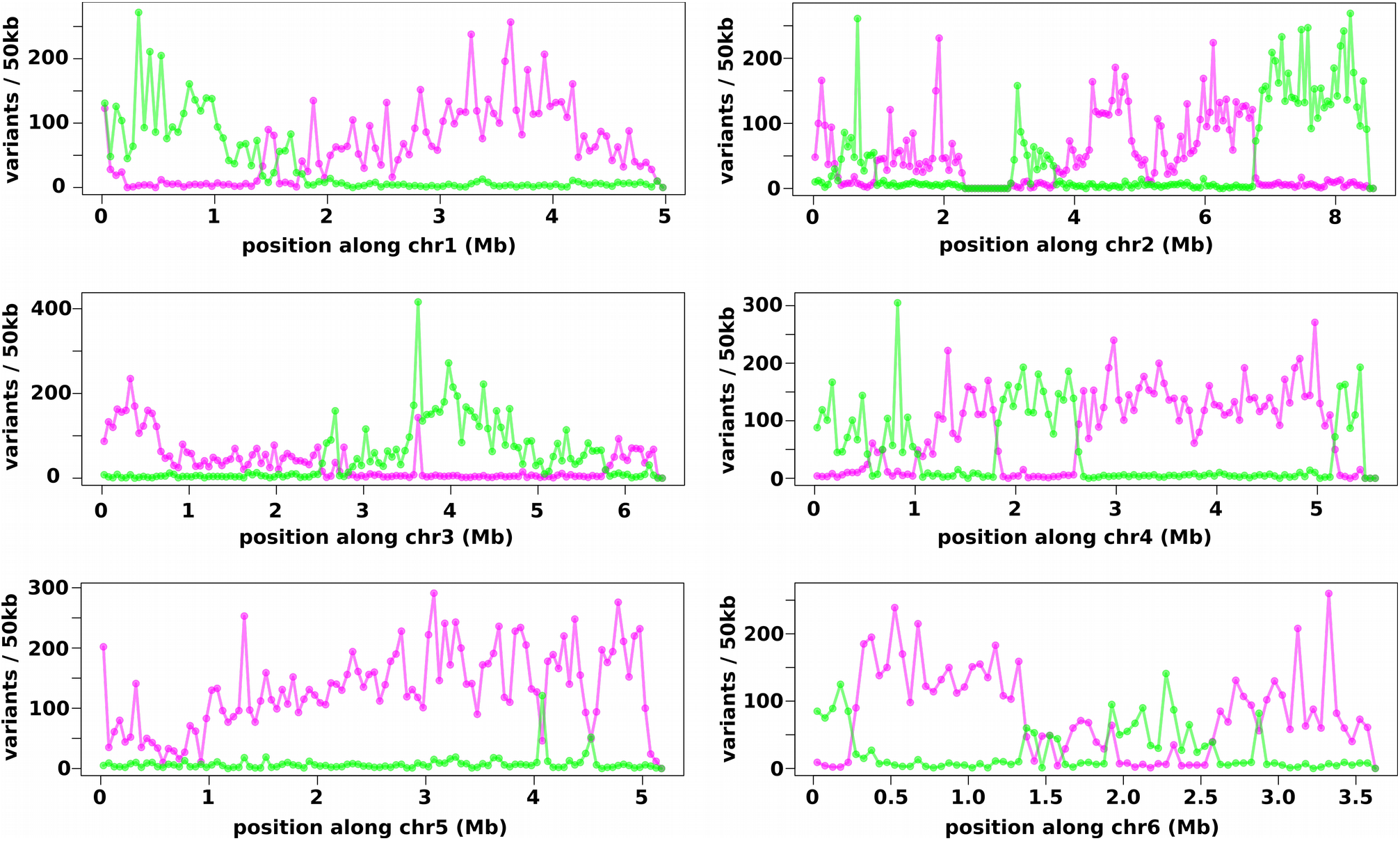
Recombination in one representative progeny clone. After whole genome sequencing of XGB1, a haploid progeny clone from the cross between HM597 and HM598, variants specific to each parent were quantified in 50 kb segments along all six nuclear chromosomes. Variants from HM597 are plotted in magenta, those from HM598 are plotted in green. *D. discoideum* chromosomes are telocentric, and centromeres are plotted to the left hand side.

**Figure 3.**
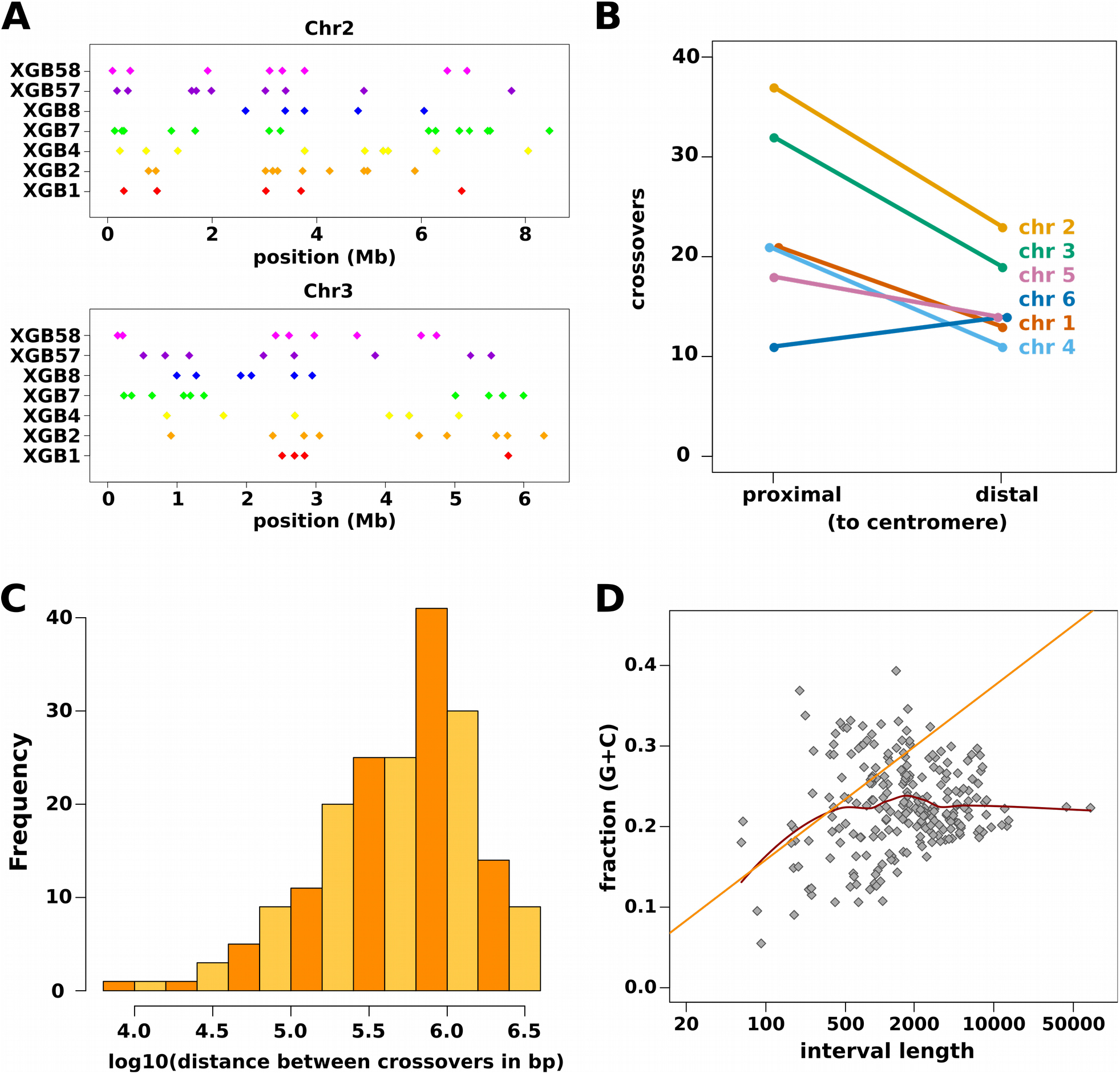
Overall patterns of recombination. **A.** The positions of crossovers along the two longest chromosomes (2 and 3) are displayed for all seven sequenced progeny clones. **B.** In five of six chromosomes there are more crossovers overall in the centromere-proximal than the centromere-distal half. **C.** Crossovers tend to be spaced hundreds of kilobases apart, although a small fraction cluster more closely. More progeny will be required to assess confidently the nature of crossover interference in social amoebae. **D.** Plotting the G+C base composition of the nucleotide sequences between the variants flanking each crossover suggests a bias in crossover designation towards low G+C regions. The straight line represents a linear model fit on the intervals shorter than 300 bp, while the curve represents a quadratic polynomial Loess least-squares fit using all intervals.

The unusual SPO11-independent recombination during sex in social amoebae has been suggested to be connected with the (anti-)social nature of macrocyst formation in which large numbers of "victim" amoebae are consumed (20); more broadly, conflict is thought to have had pervasive roles during the evolution of meiosis (23). We hypothesized that transmission of genes from consumed cells into zygotes and thence the next generation could be an evolutionary incentive that might help to explain their participation in the process, resulting in some modification to the normal process of recombination. To test this idea, we performed three-way crosses in which the mating type I and II strains used in the crosses above were mixed with an excess of *matA* null cells. These mutant cells are able to fuse with other cells to form parasexual diploids, but are unable to contribute to form fully developed macrocysts when mixed with cells of any mating type (7) (Fig. S5). We predicted that progeny from these three way crosses might inherit the bulk of their nuclear DNA from the wild-type mating type I and II parents, but might capture some quantity of DNA from the *matA* mutant cells.

As in the two-way crosses, recombination was very frequent in progeny from a three-way cross.(Figs. 3A and S4). Across all progeny characterised in two- and three-way crosses, cross-overs occurred on average approximately once per Mb in each meiosis (genetic sizes of chromosomes range from 394 to 895 cM; the 34 Mb genome was found to be 33.5 M in size in total), comparable in frequency with budding and fission yeasts (17). *D. discoideum* chromosomes are telocentric, and crossovers occurred more often in the centromere-proximal half than the distal half in five out of six chromosomes (Fig. 3B). The single chromosome in which this pattern was not observed, chromosome six, is the shortest in this species suggesting that physical distance from the centromere affects the likelihood of crossover formation. Crossovers were often regularly spaced 500 to 1000 kb apart, but were sometimes very closely spaced, within 50 kb of each other, suggesting that an interference-independent pathway operates in this species (Fig. 3C). Where flanking variants were near enough that crossover sites could be located with some accuracy, they could mostly be mapped within non-coding sequence, with a bias towards more A/T-rich sequences (Fig. 3D and Table S2). Ribosomal DNA, which is maintained in growing cells as multiple copies of a linear extrachromosomal element, was inherited largely uniparentally (Fig. S6). Additionally, one of the parental strains, WS2162, carries a high-copy nuclear plasmid that is stably carried by haploid cells during proliferation and asexual development (24); all progeny from crosses involving this strain lacked detectable plasmid as assessed by whole genome sequencing and PCR (Fig. S7).

This first genome-wide analysis of meiotic progeny in *Dictyostelium* confirms that high frequency meiotic recombination occurs in social amoebae in the absence of SPO11. Our data suggest that crossover designation is non-random, but the unconventional mechanism that has likely evolved in social amoebae remains mysterious. We have no strong candidate inducer of DNA breaks, and cannot rule out the possibility that spontaneous lesions are used to initiate recombination (20).

Returning to the question of whether lateral transmission of genes might occur from “victims” into the zygote, in progeny analysed so far from three-way crosses, we have not been able to identify any nuclear chromosomal variants that are unique to the *matA* null parent, indicating that the frequency of lateral inheritance of nuclear haplotypes from “victims” is low in *D. discoideum*, if it occurs at all (Figs. 4A and S4). Remarkably, however, in three out of four sequenced progeny from three-way crosses we found that mitochondrial genomes were inherited from the *matA* null cells (Fig. 4B). These haploid progeny have three genetic parents: two contributing nuclear haplotypes, and a third the mitochondrial genome (we refer to haploid parents and haploid progeny in this system because the diploid phase is reduced, only existing for a brief period before meiosis). In two of these progeny clones, almost all mtDNA had been inherited laterally, while in the third around 40% was of the *matA* null mitotype. Inheritance of the mitochondrial genome is therefore not strictly uniparental, but often one mitotype makes up by far the largest portion; we describe these as ‘predominant mitotypes’. Polymerase chain reactions (PCRs) designed to specifically amplify the *matA* null mitotype revealed that 13 of 42 independent progeny from the three-way cross inherited mtDNA laterally (Table S1, Fig. S8). Using fluorescent proteins targeted to mitochondria and nuclei, we could visualise thorough mixing of cytoplasms in syncytia formed during three-way crosses, indicating that lateral transmission of mitochondria most likely occurs via these transient fusions early in the sexual cycle (Fig. S9). We cannot exclude the possibility that it might also occur by escape of mitochondria from cannibalised cells within food vacuoles in the zygote, but presumably much less frequently than through mixing of cytoplasm in syncytia.

**Figure 4.**
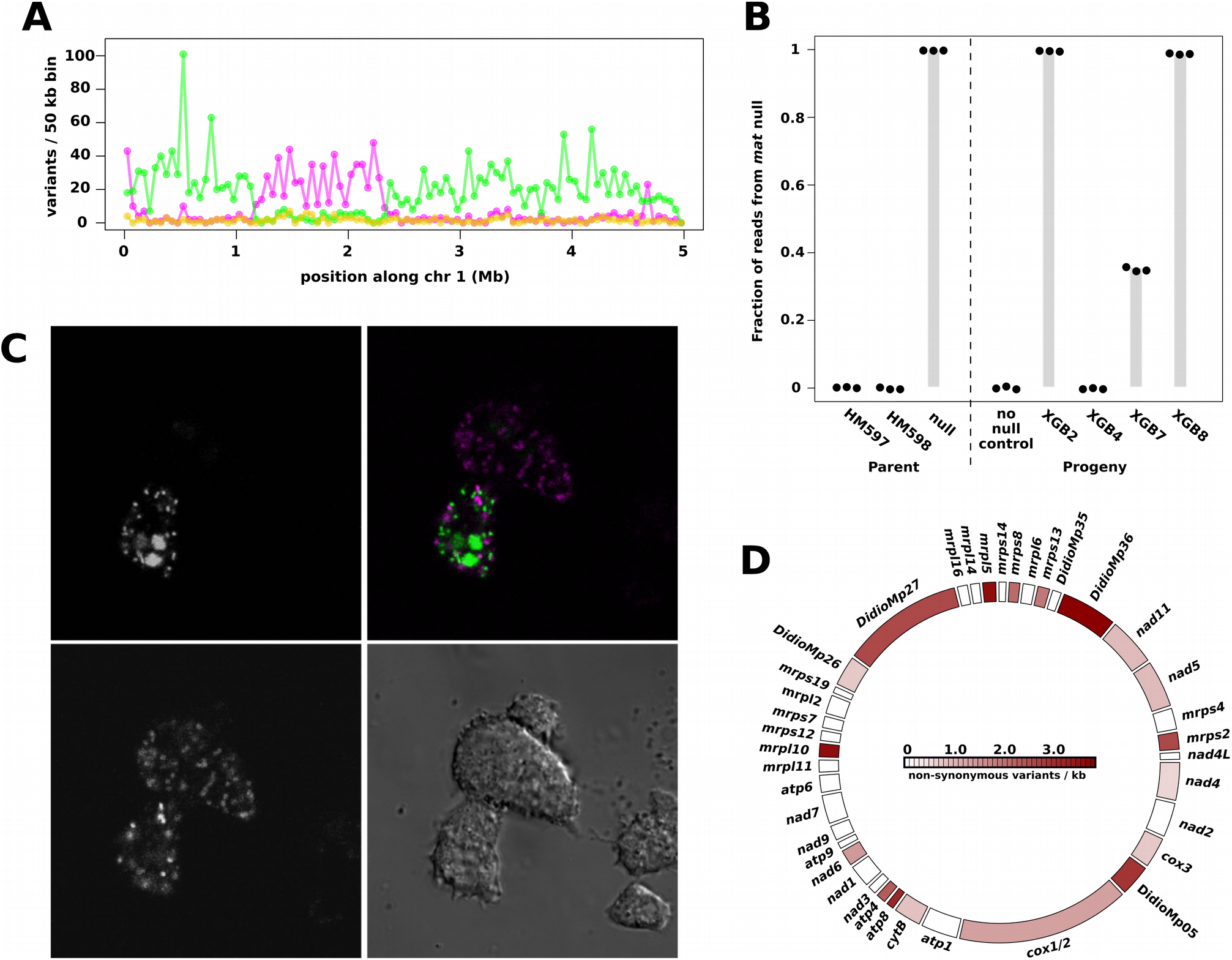
Lateral transmission of mitochondria in the *Dictyostelium* sexual cycle. **A.** We have so far found no evidence for transmission of nuclear DNA from the mutant parent in a three-way cross between HM597, HM598, and the *matA* null strain HM1524; shown here is one representative chromosome from one progeny clone, XGB8. Variants specific to HM597 are shown in magenta, HM598 variants are in green, HM1524 (AX2) variants are in gold; all putative HM1524 variants appear to be false positives. **B.** In contrast, lateral transfer of mitochondrial DNA clearly occurred in three of four sequenced progeny. Sequence reads specific to the *matA* null strain were counted at three positions along the mitochondrial genome for the three ‘parent’ strains and one two-way progeny (“no null control”, XGB1) as well as the four three-way progeny. In the three progeny with lateral transmission of mtDNA all other variants along the mitochondrial chromosome also showed similar contribution from the *matA* null. **C.** Three-way fusions were directly observed between an HM598 line expressing GFP-tagged histone 2B (green, upper left), a AX2 line expressing GFP targeted to the mitochondrial matrix with the TopA N-terminal targeting motif (green, upper left), and an HM1558 line expressing RFP anchored on the mitochondrial outer membrane using the GemA C-terminal targeting motif (magenta, lower left). A DIC image is shown lower right. Scale bar represents 10 µm. **D.** The *D. discoideum* mitochondrial genome is unevenly polymorphic: the density of non-synonymous polymorphisms in protein coding genes in the four strains used in this study are shown (homing endonuclease genes are not shown).

Our finding that one parental mitotype dominated in each progeny clone in most cases is consistent with an earlier observation in a related species (25), although in several cases we found that two mitotypes were retained in near-equal proportions (Table S1). In the two-way HM597 x HM598 cross, 10 of 11 progeny carried predominantly the HM597 mitotype and the other carried both mitotypes in near-equal quantities. For the three-way cross, the HM597 mitotype was predominant in 31 of 41 progeny, HM598 in five, the *matA* null mitotype in two, while in three progeny no single mitotype was predominant. In all five progeny in which the HM598 mitotype predominated, the *matA* null mitotype was present in small quantities (Table S1). These data indicate that mitochondrial inheritance is not strictly controlled by the mating-type locus in this species; similarly biparental inheritance often occurs in myxogastrids, one of the sister groups of social amoebae (26, 27). The over-representation of HM597 mitotypes in progeny obtained so far suggests that mitochondrial inheritance is non-random. Further crosses and backcrosses will be needed to distinguish between effects of nuclear genotype (including mating type) versus mitotypes. Mitotype inheritance frequency could simply follow passively from the overall fusion frequency of each strain. There is also clearly scope after fusion for both nuclear and mitochondrial genes to promote the inheritance of particular mitochondria.

This semi-autonomous mitochondrial behaviour in *Dictyostelium* leads us to expect that mitochondrial genes actively compete to maximise their chance of transmission to the next generation: variants that are relatively successful at promoting gamete fusion and/or subsequent survival in zygotes and transmission into haploid progeny should spread through the population, responding to a form of sexual selection. Our contrived experimental set-up involving the *matA* mutant defective in zygogenesis gives a very concrete example of how this reassortment allows mitochondria to escape from uncompetitive nuclear backgrounds, provided that cell fusion occurs. We note that because multiple mitotypes can be inherited by each *Dictyostelium* progeny clone, it is likely that some progeny have four or more genetic parents.

The asexual developmental cycle of social amoebae in which fruiting bodies are constructed with cellular stalks also engenders conflict: variants that avoid differentiating into stalk cells can act as ‘social parasites’ and spread through the population (4). We wondered whether mitochondria could spread laterally during cell fusions that occur in asexual aggregates in *Dictyostelium*. To measure this, we mixed cells containing either a fluorescent histone or mitochondrial reporter, and measured the frequency of spores containing both reporters after differentiation. These dual-labelled cells could be the result of transient fusions in which mitochondria are transferred without nuclear fusion, or more stable fusions leading to non-sexual parasexual diploid cells, which occur at a frequency of 10-6 to10-5 in similar crosses (7). We found that lateral transmission of mitochondria occurred at a frequency of around 10-4 to 10-3 (Fig. S10). This suggests, consistent with earlier reports of transient anastomoses and transfer of cytoplasm between aggregating cells (28), that transient cell fusions are more frequent than full parasexual fusions, and allow mitochondria (and other cytoplasmic elements) to be laterally transmitted. The frequency of lateral transfer in the asexual cycle found here is relatively low, but it may provide a way for self-interested cytoplasmic genomes in some strains to escape suffering the stalk cell fate, enabling dispersal and self-perpetuation. We predict that frequencies of this asexual lateral transfer vary in different isolates, likely influenced by nuclear as well as cytoplasmic genes.

Our findings suggest strong sexual selection on mitochondrial genotypes in social amoebae may occur. The four mitotypes (of the three wild-type parental strains plus the *matA* mutant) used in this study are non-uniformly polymorphic (Fig. 4D), and several of the most densely polymorphic genes are highly divergent in sequence or lacking in homology to known proteins (29), suggesting that they are evolving quickly. Most of these genes encode components of mitoribosomes and other large complexes that must be constrained in their function and evolution by their need to co-assemble with proteins encoded in the nucleus, so the extreme divergence of some of these genes is surprising. Additionally, the transcript of one poorly annotated mitochondrial gene is enriched in gametes (30), supporting the idea that polymorphic mitochondria-encoded proteins might actively influence transmission.

Sex originated early in eukaryotic evolution before the last eukaryotic common ancestor, but most likely after the first mitochondriate common ancestor (31). Before nuclear control mechanisms evolved, inheritance of mitochondria in these earliest eukaryotes must have been a result of competition between endosymbionts, and so sex would have been an opportunity for competitive mitochondrial genomes to spread in the population and escape from low quality host environments (32). One hypothesis for the origin of sex is that selfish elements first promoted cell-cell fusion as an efficient way to spread in this way (33), and it is possible that proto-mitochondria could have been the original instigators of sex. Our findings also suggest that sexual selection could be a potent force promoting the retention of certain genes within mitochondrial genomes.

We suggest that the strikingly non-Mendelian character of *Dictyostelium* sex may have consequences that help shape and perhaps resolve the conflicts engendered by the dramatically different fates of cells within groups of social amoebae. It seems likely that other important consequences will follow: the mixing of many cells’ cytoplasms during sex risks the spread of harmful endosymbionts, from selfish mitochondrial genomes to viruses and the bacteria present in certain isolates (34, 35). To the extent that out-breeding occurs in social amoebae, we expect that defences against cytoplasmic parasites must arise concomitantly.

## Materials and Methods

### Growth of *Dictyostelium* cells, and cell fusion

*Dictyostelium* cells were grown in association with *Klebsiella pneumoniae* on SM agar plates (Formedium). Fusion-competent cells were prepared by growth in shaken suspension (180 rpm) in heat-killed *K. pneumoniae* (OD_600_ of 10) in MSS buffer (5 mM MES, 10 mM NaCl, 10 mM KCl, 10 mM CaCl_2_, pH 6.0) to a density of 3-6 x 10^6^ cells per ml in the dark at 22 °C, then incubated on ice for one hour, then cleared of bacteria by differential centrifugation (3 minutes, 300 *g*, 4 °C) and washing in ice-cold MSS, repeated three times, resuspending the cells at a final density of 5 x 10^6^ cells per ml in ice-cold MSS. Cell strains were then mixed and shaken at 180 rpm and 22 °C in conical flasks to initiate fusion. Cells were imaged in Lab-tek II chamber slides (Nunc) using laser-scanning confocal microscopes (Zeiss).

### Macrocyst formation and germination

The three pairwise crosses (two-way crosses) were between HM597 (a clone made in our laboratory of the David Francis stock of A2cyc^r^, mating type II), HM598 (a clone made in our laboratory of the David Francis stock of of WS205, mating type I), and WS2162 (obtaining from the Dicty Stock Center (*36*), mating type III). A2cyc^r^ and WS205 were previously found to germinate at adequate frequencies, and to lack obvious chromosomal rearrangements (*37*) (*16*) (*38*). Macrocysts were produced by spreading 10^4^ spores of each strain to be crossed along with stationary-phase *K. pneumoniae* on LP agar plates (1 g/L peptone, 1 g/L lactose, 12 g/L agar), overlaying the cells with 10 ml MSS buffer, followed by incubation in the dark at 22 °C. In the three-way cross, 10^3^ spores of HM597 and HM598 were mixed with 1.8×10^4^ spores of HM1524, a *matA* deletion mutant. After 5-6 weeks, macro-cysts were germinated according to the method of Wallace and Raper (*37*). Briefly, macrocysts were washed from the surface of the agar plate into centrifuge tubes then centrifuged and washed three times in MSS buffer (3 minutes, 300 *g*), resuspended in 10 ml 0.005% SDS and shaken for 210 minutes at 180 rpm at 22 °C. The macrocysts were then washed twice in MSS before being resuspended in 10 ml 25 mM EDTA (pH 7.4) and shaken for 150 minutes as above. They were then washed twice in 0.025% dihydrostreptomycin solution then plated on 0.025% dihydrostreptomycin agar plates and incubated in ambient light at room temperature. Germination occurred 10-20 days later, and fruiting bodies typically emerged from within, or adjacent to, macrocyst walls. Spores were picked and cloned on SM agar plates. Putative progeny were initially checked for their *cycA* genotype by growth on SM agar plates containing 500 µg/ml cycloheximide, and screened by PCR with primers specific to *matA, matC,* and *matS* to ascertain mating type. Recombinant progeny were obtained in multiple independent experiments, although germination frequency was always low (< 1%).

### Whole genome sequencing, read mapping and variant calling

Total genomic DNA was prepared using Zymo midi X kits and used to prepare libraries using the TruSeq, TruSeq No-PCR, or Nextera Mate-Pair protocols (Illumina), followed by sequencing on Illumina HiSeq and MiSeq systems. Progeny clones were sequenced to a median depth of 25x to 149x. Genome sequences for each parent (HM597, HM598, WS2162, and AX2; HM1524 is a clonally derived mutant of AX2 and so expected to bear a haplotype near-identical to AX2) were prepared by first aligning sequence reads from each strain onto the AX4 reference sequence (*39*) (*40*) using bwa-mem (*41*) and variant-calling using Freebayes (*42*), followed by iterative consensus-building using vcftools (*43*), and remapping of reads. For strains HM598 and WS2162, additional sequence reads generously provided by Jason Wolf were included in the mapping-consensus procedure. Reads from each progeny clone (XGB1, XGB2, XGB4, XGB7, XGB8, XGB57, and XGB58) were aligned and variants called against the sequence of the parent not used in that particular cross; potential PCR duplicates were removed before variant-calling. One copy of the chromosome 2 duplication present in AX4 was masked. Variants were filtered by rejecting positions with a read depth of less than four and/or a variant quality score less than 100.

### Identification of crossover sites

A program, Hycco, was written that uses Hidden Markov Models (HMM) to identify crossovers in haploid meiotic progeny (https://github.com/tjs23/hycco) using single nucleotide polymorphisms (SNPs) listed in pairs of VCF files (*43*) derived from the three genotypes (progeny clone and two parents). Hycco works by first segmenting each chromosome into binned regions of a specified length (here we used 10 kb). Within these regions the presence of SNPs, that can distinguish one parental genotype from the other, is input to train a Gaussian HMM using the Baum-Welch method. This gives maximum likelihood estimate of the probabilistic HMM parameters using only the observable SNP data. The HMM is then interrogated for the probabilities of its underlying (hidden) parental genotype states (labelled as A or B) using the Forward-Backward method. This assigns one of the two parental genotype states (A or B) to contiguous segments of the chromosomes. Where the genotype state of highest probability swaps between A and B a chromosomal crossover is inferred. The crossover point is then more precisely estimated, at a resolution better than the initial HMM segments. Here the closest pair of parent-specific SNPs to the HMM segment edge that swap from A to B or from B to A, in the same way as the HMM state change, are sought. The crossover point is then estimated as halfway between these two A/B distinguishing SNPs. Crossover positions were confirmed by manually identifying the flanking variants.

### Enumeration of mitochondrial and ribosomal DNA variants

After mapping total genomic reads to the reference genome used in each cross and variant calling, we estimated relative proportions of each mitotype in each clone by extracting all reads mapped to the mitochondrial genome and counting reads matching k-mers exactly matching specific variants plus ten base pairs to either size. K-mers were counted around positions 8253, 14271, and 44740 on the mitochondrial genome, and 15868, 16156, and 27467 on the ribosomal DNA palindrome (coordinates in the AX4 reference genome). The numbers of non-synonymous variants per kilobase that differ between the four parental strains used in the study were colour-coded on a circular representation of the protein coding genes of the mitochondrial genome using the ‘circlize’ package in R (*44*).

### PCR screening

The HM1524 mitotype differs from the other by an 8 bp insertion relative to the other parental strains at position 6956. The two oligonucleotide primers: 5’ -CGTTTCTCTAAATTACATATATATATATATATATACTG-3’ (HM1524) and 5’ -CCGTTTCTCTAAATTACATATATATATACTG-3’ (other strains) were used in separate PCRs with the common primer 5’ -AGATTTTGGTGAAAAAATGGAGTTA-3’, using the cycle parameters 96 °C for 30 s, then 25 cycles of 10 s 96 °C, 15 s 59 °C, 60 s 68 °C, then 300 s 68 °C. For progeny without whole genome sequence data support, amplicons from this PCR were Sanger-sequenced and the specific match to the HM1524 at position 6175, at which the other parent strains differ, was confirmed. Dominant mitotype was checked by PCR using the primers: 5’ -GATGCTGTTAAAGCTTTCAATCCA-3’ and 5’ -ACTAGAAAGTTTACCTTACTAGAG-3’ and Sanger sequencing to check positions 16801 and 17193. The presence of Ddp5 was tested using the primers: 5’ -TGCCATCAATAAAATTGTTCTA-3’ and 5’ -CCAATGAAAACATCATTATATCTA-3’.

### Fluorescent protein reporters and transformation

Histone 2B was tagged at the C-terminus with GFP, cloned into a pDM vector containing a *hygR* gene driven by the *act14* promoter (*45*). The first 258 bp of the *topA* gene was cloned to the 5’ of the *GFP* gene in pDM340, which contains *neoR* driven by the *act8* promoter (*46*). A vector containing the mitochondrial targeting motif of GemA tagged to the C-terminus of tagRFP and *hygR* driven by the *act8* promoter (*47*) was a gift from Jason King. To image fusion of *matA* null cells, strain HM1524 was transformed with pDXA-GFP (*48*), HM1558 (a type III strain whose derivation will be described fully elsewhere) was transformed with pDXA-mRFPmars (*49*). Cells were transformed by electroporation (*50*) and selected in diluted SM broth with antibiotic-killed *K. pneumoniae* (*51*) in the case of strain HM598, or axenically in HL5 medium (AX2 and HM1558) in 100 µg/ml hygromycin or 20 µg/ml G418. Expression of the *topA-GFP* construct was induced by the addition of 10 µg/ml doxycycline during the pre-fusion growth step; the drug was not added during or after fusion.

### Flow cytometry and fuorescenceeactivated cell sorting

H2B-GFP and GemA-tagRFP cells were mixed in a ratio of 50:50 and co-developed on KK2 agar plates (16.5 mM KH_2_PO_4_, 3.9 mM K_2_HPO_4_, 2 mM MgSO_4_, 0.1 mM CaCl_2_, 12 g/L agar). As controls non-mixed cells were used. After at least 24 h of development the ripe spore heads were harvested with an inoculation loop and transferred into KK2 buffer. To remove the sorocarp and avoid clumping of spores the suspension was filtered using a 10 µm filter (Symex) to filter the sample. The filtrate was transferred into conical centrifuge tubes at low density (about 2×10^5^ spores per ml) and used either for cell sorting or flow cytometry. For flow cytometry the samples were run on a LSR II flow cytometer at low speed to avoid doublet events. At least 50000 spores were analysed per sample. To filter for GFP fluorescence a B525 filter and for tagRFP a YG610 filter was used.

## Acknowledgments

We wish to thank Einav Gross, Vitaliy Kuznetsov, Shun Adachi, Anastasios Tsaousis, and Robert R. Kay for helpful discussions, and Jason King and Jason Wolf for sharing reagents and data. We are grateful to the MRC EASIH and CRUK Cambridge Research Institute genomics facilities for carrying out DNA sequencing.

## Author contributions

GB conceived the project, performed crosses, germinated the macrocysts and characterized progeny, and drafted the manuscript; PP performed flow cytometry and fluorescence activated cell sorting; MO and HU tested for cell fusion and macrocyst formation; TJS wrote analysis software.

## Data and materials availability

Sequencing data and consensus parental genomes are available under the accession PRJEB28008.

